# Inference of Cichlid Speciation Patterns is Dependent on Spatial Covariance and Species Delimitation

**DOI:** 10.64898/2026.02.12.705618

**Authors:** Eleanor M. Hay, Samuel R. Borstein, Matthew D. McGee

## Abstract

Macroevolutionary analyses typically treat species as discrete units and account for shared evolutionary history. However, speciation is a continuous process and taxa are often spatially clustered, potentially biasing inferences of diversification. Here, we investigate how species delimitation and spatial non-independence influence speciation dynamics and inferred drivers using cichlid fishes as a model system. Using a phylogeny and trait dataset of 1,712 species, we first generated a reduced dataset of 820 species by removing incipient species based on known breeding compatibilities. We then fit phylogenetic and spatiophylogenetic models using an integrated nested Laplace approximation framework to jointly account for phylogenetic and spatial covariance. We find that the treatment of incipient species and spatial non-independence both alter speciation patterns and inferred drivers. Analyses of the full phylogeny identified strong trait associations and spatial hotspots driven by young adaptive radiations in Lake Victoria and Lake Malawi, whereas removing incipient species and accounting for spatial non-independence reduced extreme speciation rates, weakened or removed trait effects, and largely eliminated spatial hotspots. These results demonstrate that macroevolutionary inference is sensitive to species delimitation and spatial structure, highlighting the need to consider the influence of incipient species and spatial covariance in comparative analyses.

## INTRODUCTION

A central goal in evolutionary biology is to understand the factors responsible for generating and maintaining biodiversity. To determine these factors, a multitude of approaches have been developed to estimate speciation and extinction rates (e.g., Nee 2006; Jetz et al. 2012; Rabosky et al. 2014; Maliet et al. 2019) and examine correlations between traits and lineage diversification rates (e.g., FitzJohn 2010; Beaulieu and O’Meara 2016; Rabosky and Huang 2016). However, phylogenetic comparative methods do not come without limitations, and past studies have assessed the accuracy of different estimation methods (Title and Rabosky 2019), evaluated the validity of models (Rabosky and Goldberg 2015), and questioned the assumptions that can be made from extant time trees (Louca and Pennell 2020). It is essential to understand what we are estimating and how we are estimating it, and beyond methodological concerns, other external factors can significantly impact the outcomes of speciation methods.

Biologists have delineated biodiversity into discrete units - species - for centuries (Linnaeus 1753; Darwin 1859; Mayr 1942). Despite species being fundamental units of evolution, the question of how to define a species remains subject to ongoing debate (Hey 2006; Hausdorf 2011; Wang et al. 2020). With over 30 different species concepts in circulation (Zachos 2016), the biological species concept – which classifies individuals that can interbreed and produce fertile offspring under natural conditions as belonging to the same species (Mayr 1942) – remains the most widely recognized and accepted definition within much of evolutionary biology.

One key challenge with species delimitation is that speciation is not always a discrete outcome due to the variety of isolating mechanisms. Speciation biologists often invoke the idea of a speciation continuum (Hendry et al. 2009; Ottenburghs 2018; Stankowski and Ravinet 2021), where one side of the continuum involves species with no intrinsic incompatibilities shared with close relatives, often referred to as ‘incipient species’. Incipient species are capable of producing viable hybrid offspring, with factors such as environmentally induced mating periods, or mismatched trait combinations in hybrids preventing admixture in natural settings (Stebbins 1950; Matsubayashi et al. 2010; Baack et al. 2015; Chhina et al. 2022). On the other end of the continuum are so-called “good species” that are associated with intrinsic barriers that prevent the formation of viable offspring, often associated with traits such as incompatible genital morphology, improper gamete recognition, biased hybrid sex ratios, or a variety of hybrid traits associated with sterility or lethality (Dufour 1884; Haldane 1922; Stebbins 1950; Orr 1995; Sota and Kubota 1998; Masly 2012; Anderson and Langerhans 2015).

Sometimes, reproductive isolation can be studied in so-called hybrid zones, where the species ranges overlap and varying degrees of hybridization occur (Bigelow 1965; Hewitt 1988; Barton and Hewitt 1985). However, many closely related species currently occupy fully disjunct ranges with no direct opportunity for hybridization (Jordan 1908; Lessios 2008; Reichard et al. 2017). One classic method for overcoming these issues involves hybridization experiments to study reproductive isolation under controlled conditions (Rice and Hostert 1993; Bolnick et al. 2008; Arrieta et al. 2013). These experiments are often combinatorial across a clade of related species, which can pose challenges as the number of experiments scales exponentially with the number of species. Alternatively, experiments can be performed on pairs of closely related species (Brill et al. 2016; Feller et al. 2020), assuming a reliable phylogeny for the clade exists. Despite their clear importance to speciation research, hybridization experiments are rarely performed, particularly in animals, as they require an abundance of time, resources, and specialized husbandry expertise. Hybridization experiments are more common in plants (Grant 1981; Lexer et al. 2003), though it is still rare to observe systematic evaluation of isolating mechanisms across an entire clade.

In contrast to speciation biologists, taxonomic workers must distill the complex and often continuous nature of reproductive isolation into discrete units (Mayr 1942; Padial et al. 2010). The applied consequences of taxonomic work often require the use of morphological and genetic proxies for divergence, and as such, most taxonomic workers eschew the biological species concept as a practical means of delimitation in favor of morphological or phylogenetic species concepts, which rely on phenotypic and phylogenetic distinctiveness, respectively (Sokal and Crovello 1970; Donoghue 1985). The most common modern taxonomic workflow involves identification of unique combinations of phenotypic characters, supplemented by a small amount of molecular sequencing, typically so-called ‘barcoding’ genes (Hebert et al. 2003; Kress et al. 2005; Hollingsworth et al. 2009; Schoch et al. 2012). Unlike traditional speciation researchers, taxonomic workers will also frequently deal with rare species known from only a small number of specimens (Lim et al. 2012), or species from habitats that are difficult to sample and where field observation and/or laboratory experimentation is essentially infeasible, such as remote islands or the deep ocean (Pawar 2003; Troudet et al. 2017; Frutos et al. 2022).

The intersection between taxonomy and the biological species concepts presents particular challenges in the earliest stages of the speciation continuum, where young species have evolved some genotype and phenotypic differentiation without strong intrinsic isolating mechanisms such as gamete incompatibility (De Caprona and Fritzsch 1983; De Caprona 1986; Knight et al. 1998; Stelkens et al. 2010; Bertel et al. 2016). Incipient species whose ranges exhibit high overlap but do not collapse into hybrid swarms often represent good biological species, and systems displaying these properties are often popular models in evolutionary ecology (Kocher 2004). In the extreme case, fully sympatric species flocks, such as many African cichlid lake radiations, offer particularly dynamic examples of speciation, with some of the highest observed speciation rates in vertebrates (McGee et al. 2020). However, comparisons between old and young cichlid species flocks suggest that a relatively small proportion of young species persist to become clearly divergent lineages later in the radiation’s history (Wagner et al. 2014). Additionally, modern and paleontological data suggest that incipient species flocks can be affected by ‘reverse speciation’ collapsing incipient species into panmictic populations if environmental factors facilitating reproductive isolation, such as water clarity, suddenly change (Seehausen 2006b; Nagai et al. 2011).

Young allopatric species pose more substantial problems. As these species lack direct contact zones, researchers cannot always assess whether populations would be capable of interbreeding if spatial conditions changed to allow for gene flow, therefore necessitating this type of work to be performed in a controlled lab setting (Mayden 2002). Unfortunately, this situation is quite common, as in Australian *Mogurnda* gudgeons, which exist as young allopatric populations that are currently treated as an extreme speciation event within the eleotrid fishes (Thacker et al. 2023), though many other related lineages may not have received the attention necessary for delimitation of young allopatric populations. Similar patterns exist in some Malawi mbuna cichlids, where many ecologically divergent species exist in sympatry across rocky reefs within the lake (Konings et al. 2016), but single ecologically similar clades rarely have more than one species present at a given site. In recent years, taxonomists have begun to describe many of these allopatric populations as separate species (Stauffer et al 1997), making it difficult to assess which populations would merge if the opportunity presented itself, given their near-identical ecologies and sometimes similar color patterns. Conversely, the Lake Victoria haplochromine cichlid radiation exhibits a relatively low number of formally described allopatric close relatives, with the possible exception of *Pundamilia igneopinnis*, a melanistic sister species of *P. nyererei* that exists in rocky reefs nearby but never in full sympatry (Seehausen 1997; Selz et al. 2016; Svensson et al. 2024).

Other problems arise when researchers do not formally describe species known to behave as good biological species. Sometimes, this pattern occurs due to backlogs in species descriptions, but it can also occur when taxonomic traditions do not favor formal descriptions of incipient species, sometimes biasing results for exceptionally dynamic clades of particular importance to speciation biology (Kottelat 1997; Kottelat 1998; Minelli 2022). The threespine stickleback superspecies complex (Bell 1976; Baker et al. 2008), currently treated taxonomically as the species *Gasterosteus aculeatus*, contains many reproductively isolated incipient species, most famously the benthic-limnetic species pairs of British Columbia, as well as an extremely high number of lake-stream pairs throughout the Northern Hemisphere. However, because many stickleback traits involve the re-use of ancient genetic variants much older than their existing populations (Marques et al. 2016), establishing unique synapomorphies for each putative species has proved challenging enough that taxonomists have synonymized over 40 described species under *G. aculeatus* (Wootton 2009). Taxonomic traditions can also result in different standards for different clades; researchers use far more restrictive criteria to delimit birds than they do for other taxonomic groups, including amphibians, reptiles, and mammals (Watson 2005). It is also common for different taxonomic authorities to produce conflicting classifications; some organisms may be classified as a species according to one list, yet recognized only as a subspecies according to another list (Isaac et al. 2004).

The inconsistencies and challenges associated with reducing a complex speciation continuum into binary classifications are unavoidable, and they do have the potential to affect conservation and biodiversity estimates (Agapow et al. 2004; Mace 2004; Frankham et al. 2012). When it comes to biodiversity estimates, delimitation methods can over-split species, especially when researchers consider only a few approaches (Carstens et al. 2013) and taxa are geographically widespread and morphologically distinct (Chambers and Hillis 2020). Conversely, lumping cryptic species can underestimate total species estimates (Adams et al. 2014; Li and Wiens 2023).

While taxonomists are well aware of morphological and phylogenetic approaches to delimitation, and speciation biologists are well aware of the biological species concept, workers in phylogenetic comparative methods often occupy an unusual middle ground. Typically, phylogenetic comparative analyses are performed at a species level, implicitly relying on the morphological and phylogenetic delimitation methods utilized by taxonomists. While this approach has its advantages, particularly by utilizing natural history information from experts with extensive knowledge of a particular clade, its consequences warrant careful attention, especially when phylogenetic analyses commonly invoke traits and patterns associated with the biological species concept. From a macroevolutionary perspective, discrepancies in how scientists classify species could easily impact inferences of evolutionary patterns across clades. Many studies investigated the influence of sampling fractions or phylogenetic completeness on comparative methods (Höhna et al. 2011; Sun et al. 2020; Chang et al. 2020; Warnock et al. 2020; Mynard et al. 2023). However, researchers typically do not give species delimitation methods the same attention, even though the choice of species concept can potentially impact results.

A further, and often overlooked, issue is that incipient species often occur in relatively close geographic proximity. This spatial non-independence may cause issues when trying to investigate patterns of diversification in the presence of processes known to be major biodiversity generators, such as adaptive radiation. Adaptive radiations are often geographically restricted, typically occurring in areas that are geographically isolated with ecological opportunity, from which ecological and lineage divergence occurs from a common colonizing ancestor (Simpson 1953; Losos and Ricklefs 2009; Stroud and Losos 2016; Gillespie et al. 2020). Moreover, adaptive radiations often contain incipient species, making taxonomic boundaries difficult, which may lead to over splitting and taxonomic inflation of recognized species, further exacerbating spatial non-independence (Isaac et al. 2004; Burriel-Carranza et al. 2023). If a radiation occurs in a discrete region, variables unique to that region may spuriously appear correlated with variation in speciation rates. While these issues may affect diversification inferences on a single clade, the issue may be further heightened when investigating diversification patterns broadly, such as when multiple clades undergo adaptive radiations in close proximity (Wagner and Funk 1995; Ronco et al. 2021). While phylogenetic comparative methods would treat these clades as independent, they are spatially non-independent, which may bias results. Incorporating spatial non-independence into comparative analyses alongside phylogenetic non-independence therefore represents a potential approach for reducing bias.

Despite the importance of spatial processes in speciation, workers in comparative phylogenetic methods rarely consider the influence of spatial covariance. Geography plays a central role in shaping speciation: ecological opportunity, geographic range size, and heterogeneous environments are established drivers of lineage divergence, and allopatric speciation remains the most common mode of speciation (Coyne and Orr 2004; Hernández-Hernández et al. 2021). It is well established that species-level variables are expected to be more similar based on shared evolutionary history (Felsenstein 1985; Harvey and Pagel 1991), but species occupying the same geographic location can also exhibit similarity, particularly if traits such as temperature, latitude, and elevation are examined (Willig et al. 2003; Olson et al. 2009; Rahbek et al. 2019). The field of ecology has long recognized the potential spatial non-independence of traits and environmental variables (Kühn 2006; Legendre 1993), and several approaches for integrating spatial effects into phylogenetic analyses have been proposed (e.g., Freckleton and Jetz 2009; Dinnage et al. 2020). Nonetheless, such methods are not widely implemented and most macroevolutionary studies using phylogenetic comparative methods tend to focus on analyses that consider evolutionary covariance between species while overlooking covariance attributed to spatial position.

This paper focuses on two factors that have the potential to influence diversification estimates: firstly, how species are classified and how this might influence phylogenetic comparative methods and approaches; and secondly, the role of spatial covariance in shaping speciation patterns. To achieve this, we use cichlids as a model system to test the influence of species delimitation and spatial covariance on speciation patterns. The family Cichlidae contains over 1700 described species and represents a model clade for the study of speciation and adaptive radiations (Seehausen 2006a). Research on cichlids has contributed to our understanding of ecological speciation and adaptive radiations at a range of scales and has advanced understanding of the influences of sexual selection (Seehausen 2000), sensory drive (Maan et al. 2006; Seehausen et al. 2008), and competition (Winkelmann et al. 2014) on speciation. Previous phylogenetic studies have identified environmental variables such as ecological opportunity, lake depth, and aridity as predictors of cichlid radiations (Wagner et al. 2012; McGee et al. 2020). However, the geographically restricted nature of many cichlid lake flocks suggests that spatial covariance could play an important but currently underappreciated role in understanding cichlid speciation.

Another unique advantage of the cichlid evolutionary model system for speciation studies is the availability of data on reproductive incompatibilities (Stelkens et al. 2010; Stelkens et al. 2015), which is not typically available for most clades due to the time and cost of such experiments. These experiments suggest that intrinsic reproductive incompatibilities begin to emerge after roughly a million years of evolution. For example, members of the young Lake Malawi mbuna cichlid clade appear nearly universally capable of reproducing and producing fertile offspring, while hybridization between mbuna and non-mbuna does not always result in full viability (Stelkens et al. 2010; Stelkens et al. 2015).

We use cichlids to evaluate how the treatment of incipient species influences macroevolutionary inferences and to test the role of spatial covariance in shaping speciation patterns. Using a previously published phylogeny, trait dataset, and occurrence records (McGee et al. 2020), we first examined how reducing the phylogeny to include only lineages with evidence of emerging reproductive isolation (Stelkens et al. 2010; Stelkens et al. 2015) affects inferred speciation patterns. This approach should reduce the disproportionate influence of the extremely species-rich, young radiations in Lake Malawi and Lake Victoria, which can bias macroevolutionary analyses due to the re-use of ancient genetic variation (McGee et al. 2020; Rabosky 2020). We then tested whether accounting for spatial covariance alters speciation estimates by using spatiophylogenetic models (Dinnage et al. 2020), which include both phylogenetic and spatial random effects. This framework combines spatial observations with species-level estimates of abiotic and biotic variables and uses integrated nested Laplace approximation (INLA) to estimate the joint posterior distribution of model parameters (Rue et al. 2009; Martins et al. 2013). This approach retains variation in the size and shape of species’ ranges and the extent of overlap among species (Dinnage et al. 2020). We use these spatiophylogenetic INLA models to test how spatial structure influences associations between speciation rates and spatially correlated environmental predictors.

## METHODS

### Phylogeny, diversification variables and occurrence records

We used a previously published dataset for 1712 species of cichlid from McGee et al. (2020). This dataset consists of a phylogeny, eleven binary variables associated with cichlid diversification, and occurrence records (McGee et al. 2020). The phylogeny includes all taxonomically valid cichlid species described before 2019 and was constructed using meristic characters, nuclear and mitochondrial genes, and clade topological constraints. To obtain a continuous, tip-level estimate of speciation rate to use in analysis, we calculated the diversification rate (DR) statistic (Jetz et al. 2012). The DR statistic has been shown to more accurately reflect speciation than net diversification (Title and Rabosky 2019). We used this tip-level measure of speciation to remain comparable with the models originally implemented by McGee et al. (2020).

The dataset of diversification predictor variables consists of eleven abiotic and biotic variables that previous studies have found to be associated with cichlid diversification and adaptive radiation (McGee et al. 2020). This comprehensive set of predictor variables includes binary classifications of the following variables: water depth (small vs large depth gradient), the presence of predators, a polygamous mating system (rather than biparental care), the presence of male ornamentation, endemism/small range size, large body size, small body size, high elevation, high latitude, high rainfall (rainforest), and low rainfall (desert). We followed these binary classifications in order for our results to remain comparable to the original study.

To consider a spatial effect in analysis, we used coordinate information for cichlids from McGee et al. (2020), which were originally used to classify environmental variables for each species across their native range. Following this study, for endemic lake species, we used a set of coordinates defining the geographic expanse of the lake and assigned species endemic to those lakes the full set of coordinates for that lake. A total of 20,292 cichlid point occurrence records were obtained to use in analysis.

### Reduced dataset

To investigate the influence of the treatment of incipient species on speciation analyses, we trimmed the cichlid phylogeny by removing species known to breed and produce fertile offspring. In cichlids, the earliest known reproductive incompatibilities emerge between Malawi mbuna and other Malawi clades (Stelkens et al. 2010; Stelkens et al. 2015). Based on these studies, we chose to collapse all clades younger than the split between mbuna and their sister group, which occurred ∼2.2 MYA. Using the *‘treeslice’* function from the R package ‘phytools’ (Revell 2012), we split the phylogeny at an age of 66.2 million years from the root (∼2.2 MYA from the present), generating 163 subtrees. From each subtree, we randomly retained one species, collapsing all subtrees <2.2 MYA into single tips, resulting in a reduced cichlid phylogeny containing 820 species.

To assign trait values to the collapsed subtrees, we used ancestral state estimation. To capture uncertainty in trait reconstructions, we generated 10 reduced datasets (Tables S1-10). Discrete character states were reconstructed on the full cichlid phylogeny using the *‘ace’* function from the R package ‘ape’, assuming an equal-rates model. For each internal node corresponding to a collapsed subtree, we used the marginal likelihoods of each state to assign trait values across 10 replicate datasets. For example, if the ancestral state at a node was estimated to be state 1 with a probability of 0.7, we assigned state 1 in 7 out of the 10 reduced datasets, and state 0 in the remaining 3. This procedure was repeated for all collapsed subtrees and applied in a randomized manner across replicates, allowing us to incorporate uncertainty in ancestral states while retaining variation across datasets. Occurrence records for the removed species were reassigned to the retained species, and duplicates were removed, resulting in a total of 17,445 point occurrence records associated with the reduced phylogeny.

Using this reduced phylogeny, we calculated tip-specific speciation rates using the DR statistic (Jetz et al. 2012; Title and Rabosky 2019) to remain comparable to the full phylogeny. The DR statistic was log transformed and used as a continuous estimate of speciation rate for the INLA model. To visualize the effect of removing incipient species on speciation rates, we visualized rates on both the full and reduced phylogenies using the *‘contMap’* function from the R package ‘phytools’ (Revell 2012).

### Spatiophylogenetic analysis

To estimate the spatial and phylogenetic structure in speciation rate of cichlids, we used spatiophylogenetic modeling (Dinnage et al. 2020). This Bayesian approach uses an Integrated-Nested Laplace Approximation (INLA) framework to estimate the joint posterior distribution of model parameters (Rue et al. 2009; Martins et al. 2013) and is an approach for latent Gaussian Markov random field models, providing a significantly faster alternative to Markov chain Monte Carlo (Rue and Martino 2007; Rue et al. 2009). INLA can simultaneously deal with phylogenetic and spatial non-independence, enabling us to incorporate fixed and random effect predictors that vary spatially with a response variable measured at the species level. This framework represents an improvement on past approaches, which have typically summarized spatial variables to a single point, by retaining variation in the size and shape of species ranges and the extent of overlap between different species (Dinnage et al. 2020). Here, we used INLA to construct a spatiophylogenetic speciation model in which we test the influence of our diversification variables on speciation rates.

All models were implemented in the R, using the package ‘INLA’ (Rue et al. 2009; Martins et al. 2013). We first considered an INLA model with just a phylogenetic effect. For the phylogenetic effect, INLA requires a phylogenetic precision matrix which is the inverse of a phylogenetic covariance matrix. Prior to inverting, the phylogenetic covariance matrix was standardized by dividing by its determinant raised to the power of 1/N_species_ (Dinnage et al. 2020). We then considered a model that incorporates both the phylogenetic random effect and a spatial random effect. To model the spatial effect across the entire landscape, we used a spatial mesh. The spatial mesh was constructed over the 20,292 cichlid point occurrence records and averaged across the spatial random fields of occurrence records for each species. This approach integrates a spatial random field across each species’ distribution, so each species contributes a single datapoint to the likelihood, reducing any bias that could arise when the number of occurrence records varies between species (see Dinnage et al. [2020] for further model explanation). We assessed a variety of spatial meshes in which we varied in the coarseness of the mesh. Three different meshes consisting of 313, 1071, and 2905 vertices were used. Model comparison based on the marginal log-likelihood, deviance information criterion (DIC), and the widely applicable Bayesian information criterion (WAIC) was used to assess the different meshes. Model fit improved markedly at the finest resolution: the spatial mesh containing 2095 vertices had the highest marginal log-likelihood and lowest WAIC (Table S11) and was used in analysis for both the full cichlid phylogeny and the reduced phylogeny. Models were also repeated over a range of phylogenetic and spatial priors to assess the robustness of results and fit was compared with the marginal log-likelihood, DIC, and WAIC. Priors were found to have little influence on results (Table S12-S13). The probable influence of effects in the models were based on whether the 95% credible intervals of the effect size overlapped with zero.

INLA models were then re-fit to each of the 10 reduced datasets (Table S14). The size of spatial mesh had little effect on model results (Table S11), therefore we used the same sized mesh as used for the full phylogeny, consisting of 2905 vertices to remain comparable. For each model, we obtained posterior samples by drawing from the marginal posterior distributions using the ‘*inla.rmarginal’* function. We generated 500,000 posterior samples per model, yielding a total of 5 million samples across the ten reduced datasets. These posterior draws were then pooled to approximate the combined posterior distribution for each parameter across all reduced datasets. From these combined posteriors, we summarized parameter estimates by reporting the 0.025, 0.50, and 0.975 posterior quantiles, which we interpret as Bayesian 95% credible intervals. We did this for both the phylogenetic model and full spatiophylogenetic model. Statistical support for each effect was assessed based on whether these credible intervals excluded zero.

Spatial patterns of cichlid speciation rates were examined using the spatial random field estimated by the INLA spatiophylogenetic models. This field represents the spatially structured component of variation in speciation rates (log DR). To estimate this, we fitted spatiophylogenetic models that included only an intercept along with the spatial and phylogenetic random effects. These intercept-only models provide spatial fields representing variation in speciation rates without covariate adjustment, serving as a baseline for comparison. The mean of the spatial random field was projected onto the study region. This produced maps highlighting areas where species tend to have higher or lower modeled speciation rates than expected from phylogeny alone. The spatial fields were plotted on the same scale for both the full and reduced datasets to visualize how removing incipient species affects these patterns. Additionally, we mapped the cichlid occurrence records used in the analysis and colored points by each species’ estimated speciation rate (log DR). This enabled us to visualize how species-level speciation rates are distributed geographically and how these distributions correspond to the spatial patterns captured by the model.

## RESULTS

We found that removing incipient species strongly altered inferred associations between traits and speciation dynamics. Using the full phylogeny, the phylogenetic INLA model showed that the presence of predators, high elevation, and aridity all had negative impacts on cichlid speciation rates (Fig. 1A; Table S15). When we removed incipient species and repeated this analysis on the reduced phylogeny, the effects of predators and elevation were removed, and aridity was reduced, but still had an effect on speciation. In comparison to the full dataset, we found that the effect of male ornamentation on speciation has strengthened to have a positive effect, however confidence intervals do slightly overlap with zero (Fig. 1B; Table S16).

**Figure 1.**
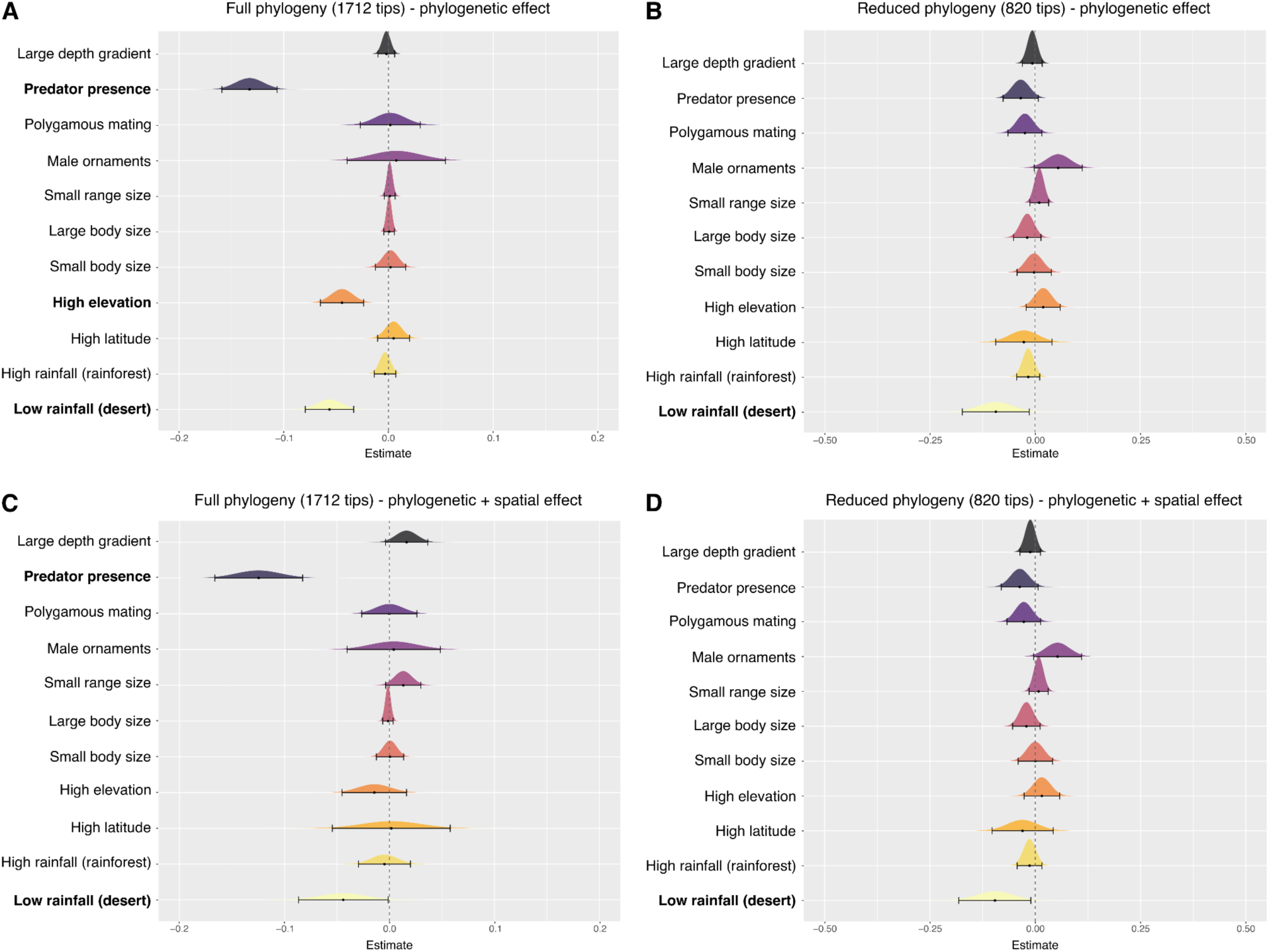
INLA model results showing the effects of fixed predictors on speciation rate. Panels display regression coefficients from the full phylogeny (A,C) and reduced phylogeny (B,D). Top panels show models with a phylogenetic effect only, while bottom panels include both spatial and phylogenetic effects. Dotted lines indicate zero, and error bars represent 95% credible intervals. Posterior distributions whose credible intervals do not overlap zero are highlighted in bold. Full model results are provided in Tables S15-16.

The inclusion of spatial covariance alongside phylogenetic covariance revealed several notable patterns. For the full phylogeny, the spatiophylogenetic INLA, which considers geographic covariance, removed the effect of elevation on speciation, and the effects of aridity and predators were reduced. Both aridity and the presence of predators still had negative impacts on cichlid speciation rates, but with weaker effect sizes and wider credible intervals (Fig. 1C; Table S15). In contrast, for the reduced phylogeny, adding a spatial random effect had little influence on parameter estimates. The results were very similar to the model with only a phylogenetic random effect: aridity negatively impacted speciation, and male ornaments had a positive effect, with confidence intervals slightly overlapping with zero (Fig. 1D; Table S16).

Spatial and phylogenetic patterns of speciation rates vary extensively in cichlids (Fig. 2). For the full phylogeny, clades containing species from Lake Victoria and Lake Malawi exhibit exceptionally high speciation rates in comparison to other lineages (Fig. 2A). This pattern is also observed in the occurrence maps, where occurrence records associated with the cichlids of Lake Malawi are associated with higher rates of speciation (Fig. 2C). Using the reduced dataset, the phylogenetic and spatial patterns change. In comparison to the full dataset, extreme speciation rates have been reduced (Fig. 2B). Occurrence records show a smoother distribution of speciation rates across Africa, with greater variation in speciation rates now visible across South America (Fig. 2D).

**Figure 2.**
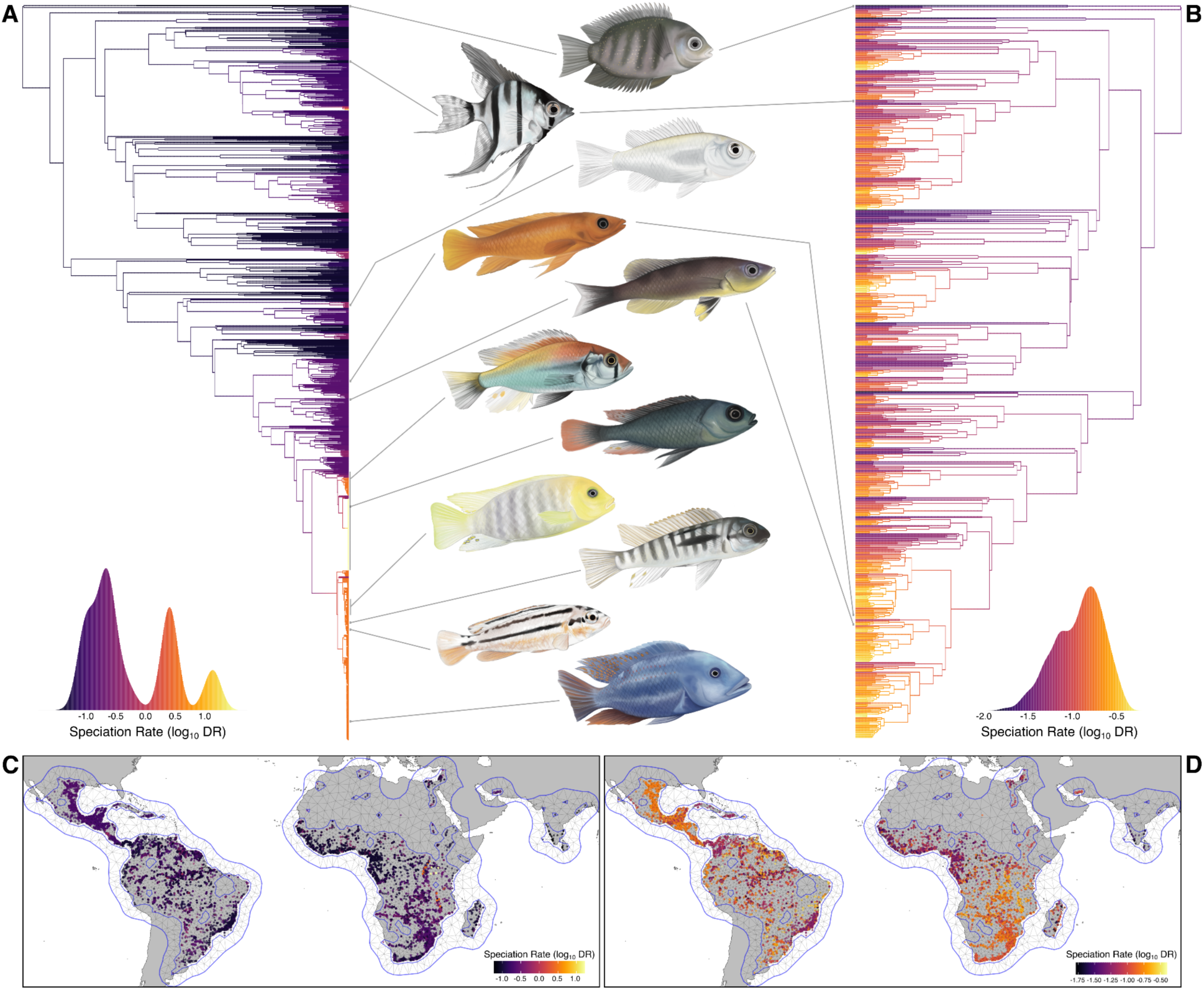
Phylogenetic and spatial patterns of cichlid speciation rates. (A) Full phylogeny of 1,712 cichlid species, with branches colored by speciation rate (log_10_DR) and the color scale displayed as a density plot beside the tree. (B) Reduced phylogeny of 820 species, with branches colored by speciation rate (log_10_DR) and color scale density plot. Maps show species occurrences for the full (C) and reduced datasets (D), colored by estimated speciation rate, with the spatial mesh used in the spatiophylogenetic model overlaid. Illustrations by Eleanor Hay.

The spatiophylogenetic models reveal pronounced spatial structure in cichlid speciation rates (Fig. 3). The mean posterior spatial random field from intercept-only models, which account for phylogenetic covariance, captures spatial variation in speciation rate (log_10_DR) that is unexplained by phylogeny. In the full dataset, regions of East Africa, particularly Lake Victoria and Lake Malawi, show higher estimated speciation rates relative to phylogenetic expectations. Most other areas show neutral spatial effects, while lower-than-average rates occur near Lake Turkana (Fig. 3A). In the pruned dataset, spatial effects around Lake Victoria and Lake Malawi are reduced because most of the species in these clades were removed. In comparison to the full dataset, spatial effects are weaker across Africa and are largely neutral across the landscape (Fig. 3B). When this spatial field is plotted on a finer scale, variation in spatial effects is apparent, with higher rates of cichlid speciation in South America and Central Africa, and lower rates of cichlid speciation in Western Africa and India (Fig. S1).

**Figure 3.**
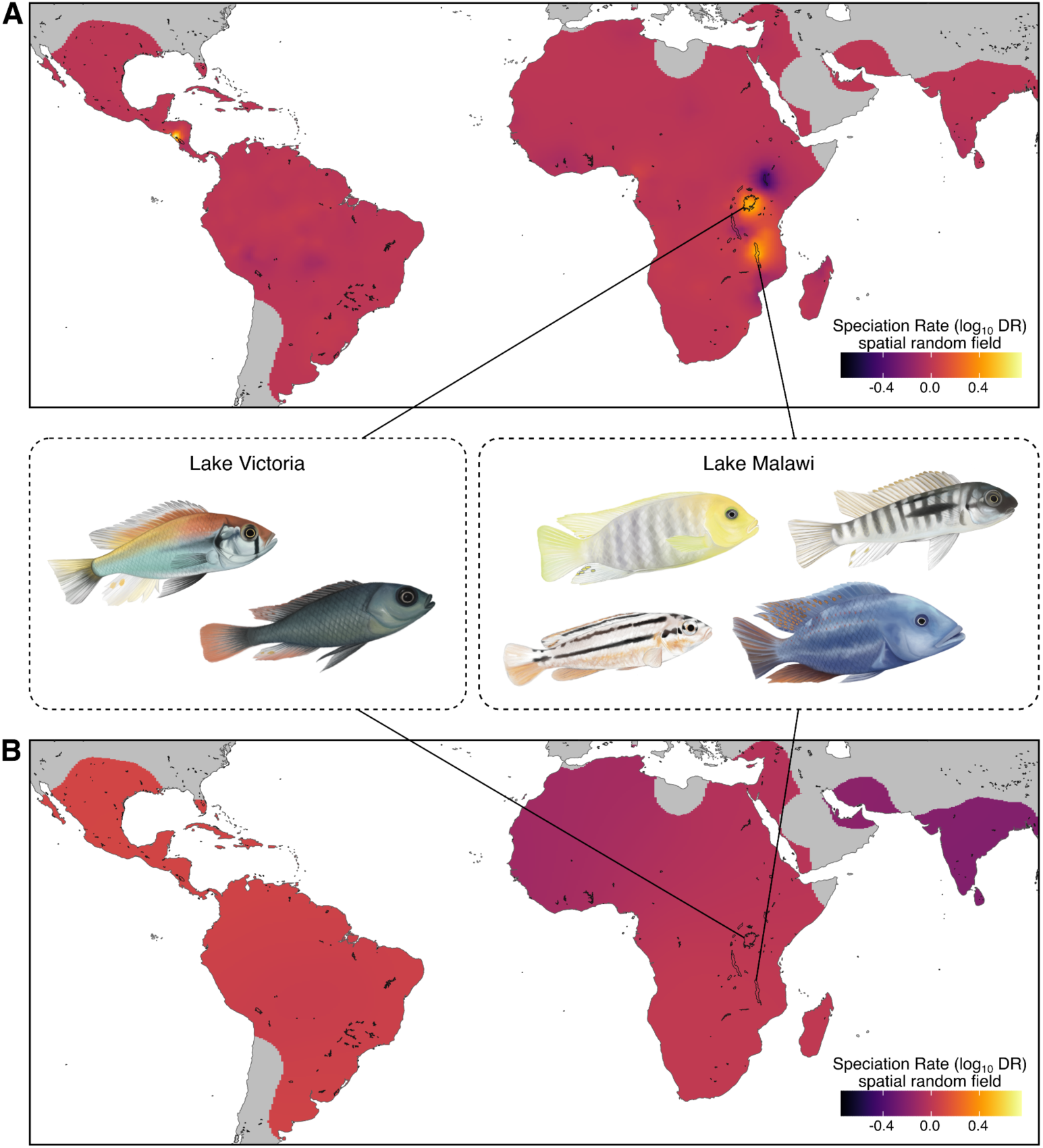
Spatial variation in cichlid speciation rates. Maps show the mean spatial random field estimates from intercept-only INLA spatiophylogenetic models, representing residual spatial structure in speciation rates (log_10_DR) after accounting phylogenetic non-independence. Panels compare the full dataset of 1,712 species (A) and the reduced dataset of 820 species (B), plotted on the same scale. Warmer colors indicate regions where species tend to have higher modeled speciation rates than expected from phylogeny alone, while cooler colors indicate regions with lower rates. Note that the mapping of rates in some regions (e.g. Florida), is due to artifacts in generating the mesh and not due to actual occurrence in the region. See Fig. 2 for occurrence records of cichlids. Illustrations by Eleanor Hay.

## DISCUSSION

Our results indicate that the treatment of incipient species exhibits a strong influence on the association between traits and speciation dynamics, with spatial covariance playing a similar role in many circumstances. While largely congruent with previous speciation analyses in cichlids, our results suggest that close attention to species delimitation does have a strong effect on downstream phylogenetic analyses. Cichlids provide what may be a worst-case scenario highlighting the necessity to consider incipient species and spatial non-independence when assessing the drivers of speciation given they display some of the fastest known rates of speciation in vertebrates (McCune 1997), much of which is concentrated in species flocks of several large young lakes (McGee et al. 2020). In such systems, failure to account for incipient species and spatial non-independence risks conflating processes associated with rapid, localized radiations with longer-term drivers of diversification.

In the original study, McGee et al. (2020) found that Bayesian regression and FiSSE analyses suggested that the presence of large visually oriented predatory fish and an arid climate are major constraints to speciation rate in cichlids, whereas male-restricted ornamentation (indicative of evolution in response to sexual selection) and a wide gradient of water depth show weak positive effects. Notably, while these results were seen in a regression analysis and with the FiSSE method, hidden-state models of speciation and extinction only uncovered an effect of aridity reducing speciation. However, this pattern was driven by a hidden state and not aridity itself, suggesting aridity has a minimal role on the observed diversification patterns of cichlids.

Our INLA analysis of the full dataset with no removal of incipient species produced similar results to the phylogenetic regression in McGee et al. (2020). When incipient species were pruned, we only observed an effect of aridity reducing speciation rates. This finding aligns with the hidden state results in the original study, which also only found evidence for aridity reducing speciation. Aridity likely constrains speciation rates across a number of cichlid tribes. For example, haplochromine and oreochromine cichlids exhibit very high species richness in lake species flocks but very low diversity in areas with desertified regions, such as *Astatotilapia desfontainii* and *Danakilia* in northern Africa (Stiassny et al. 2010; Freyhof et al. 2021), and *Astatotilapia flaviijosephi* and *Iranocichla* in the Middle East (Esmaeili et al. 2016; Çiçek et al. 2023). These patterns are consistent with aridity acting as a broad ecological constraint on speciation potential.

The pruned results largely eliminate the negative effect of visual predator presence on speciation rates. This is likely because the incipient species pruning entirely eliminates young species flocks like the Lake Victoria Region Superflock, the Barombi Mbo crater lake cichlid radiation, and some of the Central American *Amphilophus* crater lake flocks. Additionally, the pruning of incipient species reduces the Lake Malawi radiation to a small handful of representatives of major clades. Each of these young systems exhibits reduced presence of visual predators relative to surrounding riverine systems (Jackson 1961). It is therefore unsurprising that the removal of these young flocks removes the signal of predators influencing speciation rates, as remaining predator-reduced clades, such as the ancient cichlid tribes of Madagascar, do not exhibit unusually rapid rates of speciation.

Removing incipient species also strengthened the positive relationship between male ornamentation, a simple proxy for the effects of intense sexual selection, and speciation rate. While haplochromine cichlids in Lake Victoria and Malawi are known for both rapid speciation and male ornamentation, cichlids from the same tribe also possess male ornamentation and a generally higher rate of speciation, even with incipient species removed. Additionally, clades such as the South American *Apistogramma*, which exhibit both exceptionally high variation in male color patterning and elevated speciation rates, likely contribute to this relationship (Romer 2006). These results are compatible with the results of Wagner et al. (2012), which identified sexual selection as a key driver of adaptive radiation in cichlid lake endemics. Our results suggest that sexual selection may be a more potent long-term driver of increased speciation than the factors associated with the production of often evolutionarily short-lived species flocks, and highlight the value of carefully scrutinizing the treatment of incipient species in macroevolutionary analyses.

The spatial results on the full tree reveal speciation hotspots associated with Lakes Malawi and Victoria, consistent with the results of McGee et al. (2020). However, when incipient species are pruned, these speciation hotspots are removed and spatial effects are largely neutral for the reduced dataset. This is likely because a number of lineages contributing to the East African lake radiations, such as the haplochromines, tilapines, and oreochromines, exhibit high levels of diversity through much of sub-Saharan Africa. The higher speciation rates of these groups likely explains the pattern of lower speciation rates observed at higher latitudes in Africa in the spatial random field in the pruned analyses, in contrast to other studies showing increased speciation rates at high latitudes (Fig. 3). We expect that the ability of INLA methods to identify geographic hotspots of speciation rate to be of noticeable utility in future studies for many groups, especially for those composed of young species flocks occurring in close geographic proximity.

A key issue suggested by our analyses is the ability of very rapidly speciating young radiations to mask speciation patterns in other clades. Within cichlids, the exceptional species flocks within Malawi and the Lake Victoria Region Superflock exhibit such high rates of speciation that any trait-based analysis of speciation rate necessarily becomes associated with these radiations. This extreme speciation rate variation can also confound many phylogenetic methods - for example, in the original manuscript (McGee et al. 2020), many traits showed associations with speciation rate when using the simple FiSSE method, but few of these associations were validated using hidden-state models, which are better able to account for rate variation due to additional unmeasured factors and provide more complex character-independent null models of diversification.

One striking finding is that the INLA models accounting for incipient species produced nearly identical results to the model that also incorporated spatial non-independence. While adaptive radiations may occur at the continental scale, it is not uncommon for adaptive radiations to produce many species that co-occur in sympatry or are parapatric that may not have full reproductive isolation (Matsubayashi & Yamaguchi 2022). Our cichlid dataset demonstrates that pruning incipient species significantly reduces the number of species in biodiversity hotspots for cichlids, such as the African Great Lakes, which contain several adaptive radiations. Importantly, incorporating spatial covariance may capture some effects of these incipient radiations, offering a practical approach when detailed information on reproductive incompatibilities is unavailable.

While our study’s treatment of incipient species relies on experimental studies of cichlid reproductive incompatibilities (Stelkens et al. 2010; Stelkens et al. 2015), these experimental studies were limited to the largest and most diverse cichlid tribe, Haplochromini, and assumes that other cichlid tribes accumulate incompatibilities on roughly similar timeframes. Ideally, these haplochromine experiments could be corroborated via studies on other cichlid clades, though many species are likely unsuitable for such experiments due to large body sizes and aggression. Despite these issues, there are several clades of smaller species, most notably the West African *Pelvicachromis* and the South American *Apistogramma*, which would also be amenable to experimental assessment of reproductive incompatibilities.

Our results underscore several issues worth examining from the perspective of microevolutionary speciation research. Recent manuscripts have highlighted incongruities and difficulty using microevolutionary patterns to predict macroevolutionary trends (Tsuboi et al. 2024; Schluter 2024). We suggest that some of these incongruities may be associated with comparisons between the biological species concept and the species concepts utilized by taxonomists, rather than by our inability to understand the underlying mechanisms of speciation. One solution to these issues involves increasing the rate at which speciation biologists facilitate species descriptions. In classic speciation systems it can be relatively easy to describe new species because key phenotypic differences are already known, vastly simplifying the necessary taxonomic workload. Unfortunately, many reproductively isolated incipient species, such as the well-known benthic-limnetic stickleback species pairs of British Columbia, remain undescribed (Taylor & McPhail, 1999). Formally recognizing these taxa could substantially alter macroevolutionary speciation patterns within the Gasterosteidae, which currently possesses a relatively average speciation rate compared to related clades (Rabosky 2020), sharply contrasting with speciation studies in the group. Our results caution that macroevolutionary studies should take great care in invoking hypotheses associated with the biological species concept, particularly if analyses involve only traditionally taxonomically described species.

The field of ecology has long accepted how spatial non-independence may impact inference (Kühn 2006; Legendre 1993) and the topic has been acknowledged by evolutionary biologists (Freckleton and Jetz 2009; Dinnage et al. 2020), yet, spatial non-independence often remains overlooked in the phylogenetic comparative methods field. Our results demonstrate that accounting for both incipient species and spatial non-independence can improve inference of speciation patterns and the identification of potential drivers of diversification. We utilize a multiple regression approach that simultaneously considers several biotic and abiotic factors that potentially influence speciation. In contrast, phylogenetic comparative methods that incorporate state-dependent diversification typically only focus on a single trait hypothesized to drive lineage diversification (Maddison et al. 2007; Beaulieu and O’Meara 2016; Rabosky and Goldberg 2015; Herrera-Alsina et al. 2019, though see FitzJohn 2012; O’Meara et al. 2016; Nakov et al., 2019). Many of these models differ relative to our approach in that they jointly estimate discrete character evolution and various lineage diversification parameters from the phylogeny (Maddison et al. 2007; Beaulieu and O’Meara 2016). Our analyses instead rely on the DR statistic, which only reflects speciation rates (Title and Rabosky 2019) and does not attempt to estimate character evolutionary histories.

While state dependent diversification models provide an accessible way to test for the importance of a trait or character on diversification rates, understanding the drivers of evolution is complex. It is highly unlikely that a single trait fully explains observed patterns of diversification, and several hypothesized key drivers of lineage diversification have been shown to have complex histories that do not fully support state-dependent diversification (Cacho et al. 2010; Miller et al. 2021; Borstein et al. 2024). Hidden state speciation and extinction models provide an exciting and more robust avenue to test trait-dependent diversification by modelling variance in diversification that can be attributed to an unmeasured hidden state (Beaulieu and O’Meara 2016; Caetano et al. 2018), but these models currently lack the ability to account for spatial non-independence among taxa. While such a model would be a welcome addition to the field, we do not directly compare our results to those of a hidden state speciation and extinction model here. Nonetheless, such analyses could provide similar conclusions to previous studies on cichlids (McGee et al. 2020), in which hidden state speciation and extinction models largely supported character-independent diversification despite trait associations inferred from Bayesian regression models.

Our results show that macroevolutionary analyses of speciation can be strongly affected by the treatment of incipient species and ignoring spatial covariance. Comparisons between models with and without incipient species, and with and without spatial non-independence, reveal that factors associated with rapid, spatially clustered, short-lived radiations can dominate comparative analyses, masking the drivers of more persistent diversification. By showing that accounting for incipient species and spatial non-independence alters trait–speciation associations, we highlight the need for greater integration between spatially explicit approaches, species concepts, and phylogenetic comparative methods. More broadly, our findings emphasize that speciation is a spatially structured and multivariate process, and that improving macroevolutionary inference will require models that better accommodate this complexity. While it is unlikely that the long-running debate on species concepts and their effects on how we understand and manage biodiversity will be ending in the near future, we are optimistic that the increasing abundance of whole-genome and spatial data may provide more satisfying ways to address these issues.

## ACKNOWLEDGEMENTS

The manuscript benefited heavily from discussions at the 2025 Gordon Research Conference on Speciation in Ventura, CA. We would like to thank B. C. O’Meara, C. C. Nice, and N. H. Martin for providing comments on the manuscript.

